# Activation Dynamics of Ubiquitin Specific Protease 7

**DOI:** 10.1101/2023.01.11.523550

**Authors:** Gabrielle J. Valles, Nancy Jaiswal, Dmitry M. Korzhnev, Irina Bezsonova

## Abstract

Ubiquitin-specific protease 7 (USP7) is a deubiquitinating enzyme responsible for the regulation of key human oncoproteins and tumor suppressors including Mdm2 and p53, respectively. Unlike other members of the USP family of proteases, the isolated catalytic domain of USP7 adopts an enzymatically inactive conformation that has been well characterized using X-ray crystallography. The catalytic domain also samples an active conformation, which has only been captured upon USP7 substrate-binding. Here, we utilized CPMG NMR relaxation dispersion studies to observe the dynamic motions of USP7 in solution. Our results reveal that the catalytic domain of USP7 exchanges between two distinct conformations, the inactive conformation populated at 95% and the active conformation at 5%. The largest structural changes are localized within functionally important regions of the enzyme including the active site, the ubiquitin-binding fingers, and the allosteric helix of the enzyme, suggesting that USP7 can adopt its active conformation in the absence of a substrate. Furthermore, we show that the allosteric L299A activating mutation disturbs this equilibrium, slows down the exchange, and increases the residence time of USP7 in its active conformation, thus, explaining the elevated activity of the mutant. Overall, this work shows that the isolated USP7 catalytic domain pre-samples its “invisible” active conformation in solution, which may contribute to its activation mechanism.

## INTRODUCTION

Ubiquitination is a post-translational modification that can affect the localization, stability, interactions, and activity of the modified substrate. Deubiquitinating enzymes (DUBs) can reverse the effects of ubiquitination by cleaving the isopeptide bond between ubiquitin and the substrate [1-3]. The human genome encodes for nearly 100 DUBs, which are categorized into seven families [4]. About half of all DUBs belong to the cysteine ubiquitin-specific proteases (USP) family, making this the largest DUB family. Ubiquitin-specific protease 7 (USP7), also known as Herpesvirus-associated ubiquitin specific protease (HAUSP), has been extensively studied due to its involvement in a range of cellular processes and significant roles in human diseases (reviewed in [5, 6]). Most notably, it plays a role in the tumor suppressor p53 and E3-ubiquitin ligase Mdm2 signaling pathway, where USP7 deubiquitinates and subsequently stabilizes both [7].

USP7 is a 128 kDa enzyme that contains seven domains: TRAF-like, catalytic, and five ubiquitin-like (UBL) domains. All domains have been extensively characterized and have known roles in substrate recognition and enzymatic activity [5, 6]. Additionally, USP7 contains a 19-amino acid unstructured tail at its very C-terminus (residues 1084-1102) that makes essential contacts within the catalytic domain to promote its activation [8-10].

All USPs share a conserved catalytic domain that assumes a characteristic papain-like architecture [11, 12]. The USP7 catalytic domain (USP7-CD) is unique among most other USPs as it exists in a catalytically inactive conformation and requires substrate-binding for enzymatic activation [10, 13-15]. The crystal structures of the apo-USP7-CD (PDB ID: 1nb8, 4m5w, 4m5x, 2f1z, 5fwi, 5j7t)[9, 16-19] and USP7-CD in complex with ubiquitin (PDB ID: 1nbf, 5jtv, 5jtj) [9, 16] provide two snapshots of distinct active (USP7-CD_A_) and inactive (USP7-CD_I_) conformations of the protein, respectively (Figure 1A-B). In addition to the rearrangement of the active site itself, the activation switch involves significant structural changes in other functionally important regions of the catalytic domain (Figure 1C); however, the regulatory mechanism of USP7-CD activation is still poorly understood.

**Figure 1:**
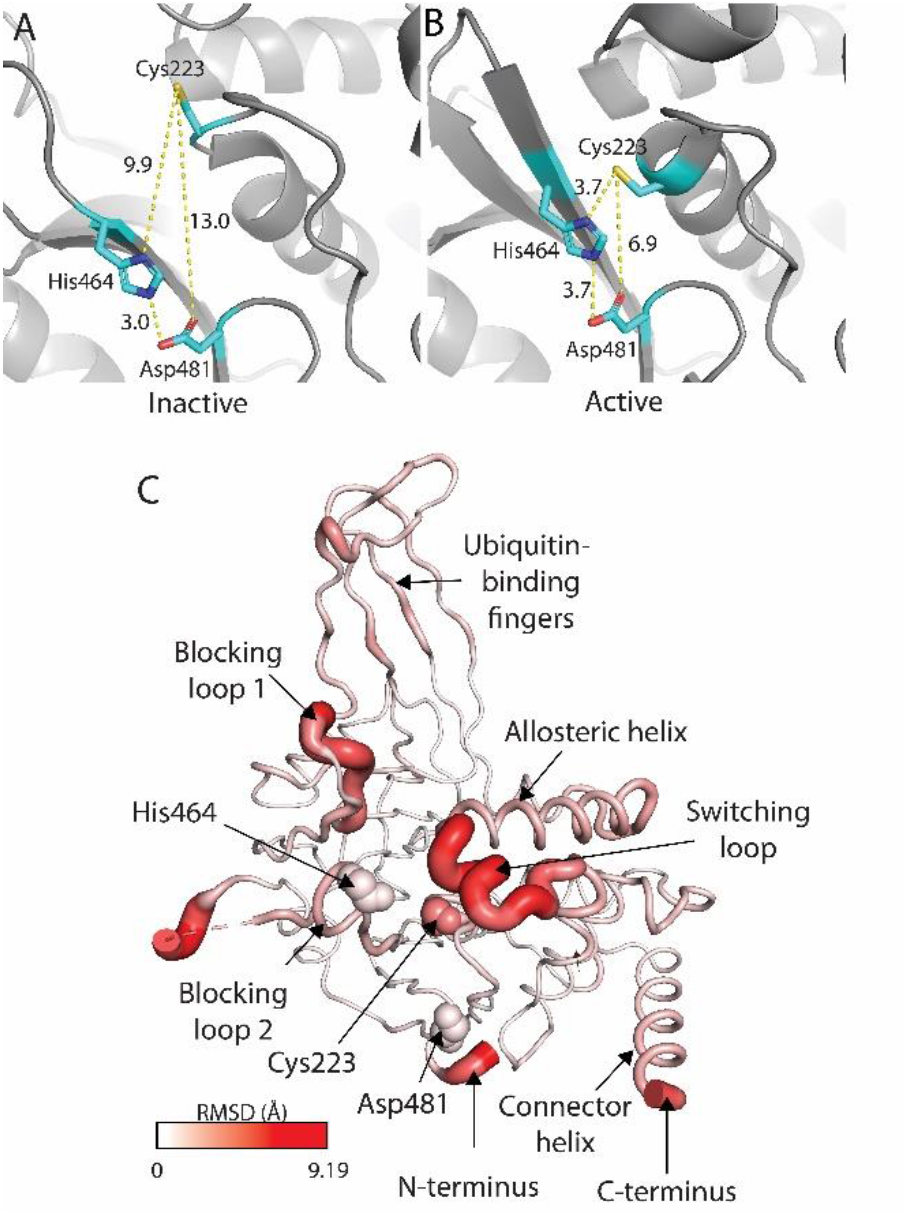
Isolated catalytic domain of USP7 can adopt distinct inactive and active conformations. (**A**) Arrangement of the active site triad residues in inactive apo-enzyme (PDB ID: 1nb8). (**B**) Rearrangement of the catalytic riad into active conformation in the presence of a substrate (PDB ID: 1nbf). (**C**) Structural changes of USP7 associated with enzyme activation. Cα RMSD between inactive and active conformations of USP7 are shown on the structure as a red color gradient.

Here, we observed the dynamics of USP7-CD using nuclear magnetic resonance (NMR) Carr-Purcell-Meiboom-Gill (CPMG) [20, 21] relaxation dispersion (RD) studies, which uniquely permit observations of conformational exchange that occurs on a μs-ms timescale, and detect low populated and transiently sampled species [22]. We report that in solution, apo-USP7-CD samples both USP7-CD_I_ and USP7-CD_A_ conformations, transiently switching between the former ground state and the latter minor state. Additionally, the largest structural movements between USP7-CD_I_ and USP7-CD_A_ localize to residues within important USP7 regions involved in activation and substrate-binding. Together, these details of USP7-CD dynamics are consistent with spontaneous enzyme activation, even in the absence of a substrate.

## RESULTS

### Isolated catalytic domain of USP7 exists predominantly in inactive conformation in solution

Overall, the USP7-CD_I_ and USP7-CD_A_ crystal structures are highly similar, with the average Cα RMSD of 1.3 Å [16]. However, key regions of the catalytic domain demonstrate dramatic conformational changes between the two structures (**Figure 1**). In the USP7-CD_I_ structure, the active site residues are positioned too far apart for the catalytic reaction to occur, with nearly 10 Ǻ distance between the nucleophile (Cys223) and the proton acceptor (His464), rendering an inactive catalytic domain (**Figure 1A**). This contrasts with the active site arrangement in the USP7-CD_A_ structure, where these residues drastically shift towards each other and the catalytic domain assumes its catalytically competent active conformation [16] (**Figure 1B**). In addition to the active site rearrangement, activation results in large structural changes within the loops surrounding the active site such as blocking loop 1 (BL1) and the switching loop (SL), as well as more distant allosteric helix and the ubiquitin-binding fingers (**Figure 1C**) [8, 16]. In the USP7-CD_I_ structure, the SL assumes an inhibitory α-helical conformation that is oriented inwards, sterically blocking the ubiquitin tail from entering the active site for cleavage (**Figure 2A**, salmon). This contrasts with the USP7-CD_A_ SL, where the loop unfolds and elongates outwards, thereby allowing ubiquitin to enter the active site for cleavage (**Figure 2A**, red) [8, 9, 16].

**Figure 2:**
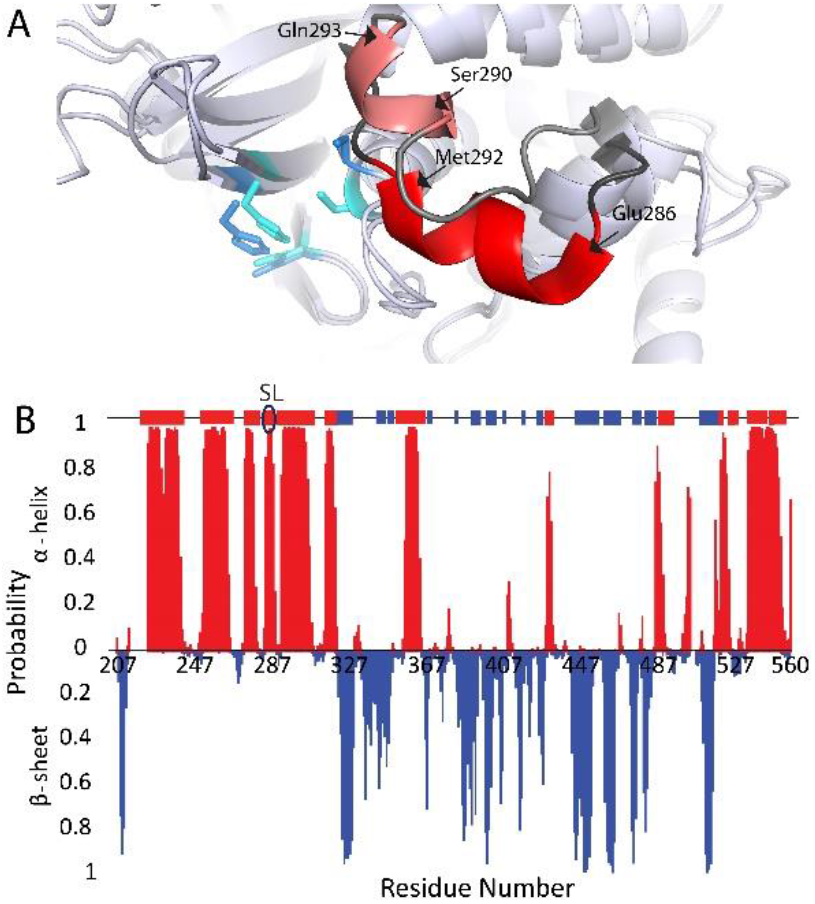
USP7 catalytic domain exists predominantly in inactive conformation in solution. (**A**) The switching loop of USP7 in inactive (salmon) and active (red) conformations of the enzyme shown in accordance with their crystal structures 4m5w and 1nbf, respectively. (**B**) NMR-based secondary structure prediction of the apo USP7 in solution. The region that corresponds to the switching loop is circled on the ribbon depiction.

We have previously reported NMR backbone chemical shift assignment of the USP7-CD [13], which are strongly correlated with protein secondary structure. Here, we used TALOS+ and experimental Cα, Cβ, CO, N and HN NMR chemical shifts to predict USP7-CD secondary structure elements in solution, (**Figure 2B**, red α-helix and blue β-strand), and then compared results to the crystal structure arrangements in USP7-CD_A_ and USP7-CD_I_ states of the enzyme. Overall, the TALOS+ predicted secondary structure arrangement matched well with USP7 structural elements (**Figure 2B**, bar graph and secondary structure elements above). Importantly, the SL (residues 283-295) was found in inhibitory α-helical conformation, indicating that USP7-CD exists in predominantly inactive conformation in solution.

### USP7 catalytic domain pre-samples enzyme’s active conformation in solution

To test whether USP7-CD can sample alternative low-populated conformations in solution we probed conformational dynamics of the enzyme using NMR CMPG RD experiments. This is a powerful and highly sensitive technique that permits the observation of protein dynamics in the μs-ms regime, which is the timescale of large global motions associated with enzyme activation. The RD approach provides detailed information about the global exchange such as the interconversion rate constant (k_ex_) and the populations of the exchanging species (p(A/B)), as well as residue-specific information reflecting local structural differences, such as NMR chemical shift differences between the species (Δω). We probed both the backbone and side chain dynamics by observing RD of ^15^N-labeled backbone amide groups and selectively ^13^C-labeled methyl groups of Ile, Leu and Val residues (ILV). The 84 ILV residues provided a good reporter coverage as they are well dispersed within the catalytic domain of USP7 (**Figure 3**).

**Figure 3:**
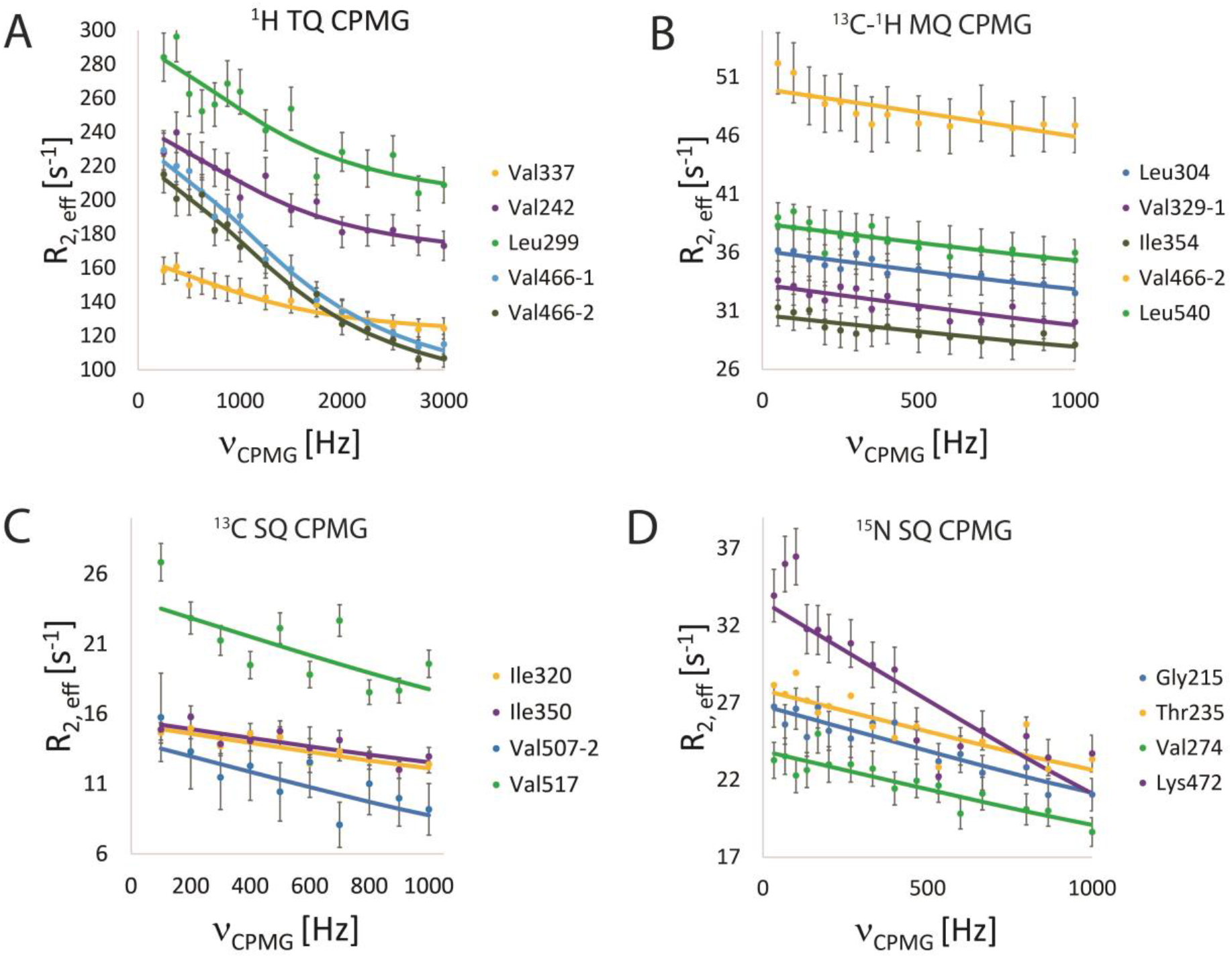
Representative CPMG relaxation dispersion profiles of the USP7 catalytic domain. (**A**) ^1^H methyl triple quantum CPMG. (**B**) ^13^C-^1^H methyl multiple quantum CPMG. (**C**) ^13^C methyl single quantum CPMG. (**D**) ^15^N amide TROSY single quantum CPMG.

A combination of ^15^N TROSY CPMG experiment for the backbone amide groups [23] and ^1^H triple-quantum (TQ) [24], ^13^C single quantum (SQ) [25] and ^13^C multiple quantum (MQ) [22] CPMG experiments for ILV ^13^CH_3_ side-chain methyl groups were performed. All RD experiments showed evidence of conformational exchange. As expected, TQ experiments provided enhanced sensitivity that resulted in higher quality data compared to SQ and MQ, thereby allowing for a more robust extraction of dynamic parameters (**Figure 3A-C**) [26]. The best quality TQ data sets were fit to a two-state exchange model to obtain the population of the minor state p(B)= 4.7 ± 0.01% and the exchange rate constant k_ex_ =7,800 ± 520 s^-1^ shown in **Table 1**. Assuming the p(B) and k_ex_ to be constant global parameters allowed us to extract per-residue Δω for ^15^N amide and ILV methyl ^1^H and ^13^C nuclei from the remaining CPMG experiments. The resulting Δω values identified specific regions of USP7 with largest structural differences between the major and minor exchanging conformations. **Figure 4** shows the location of the residues involved in the exchange color-coded from white (no exchange) to blue (largest conformational change) as well as the bar graphs of Δω extracted from individual CPMG experiments.

**Table 1:** Parameters of conformational exchange of USP7 CD and its mutants.

| | Active state population ($p_{(B)}$), % | Exchange rate constant ($k_{ex}$), $s^{-1}$ | Residence time ($1/k_{ex}$), $\mu s$ |
| --- | --- | --- | --- |
| WT | $4.7 \pm 0.01$ | $7,800 \pm 520$ | $128 \pm 8$ |
| C223A | $5.4 \pm 0.6$ | $8,984 \pm 1,112$ | $111 \pm 14$ |
| L299A | $3.5 \pm 0.2$ | $4,551 \pm 773$ | $220 \pm 30$ |

**Figure 4:**
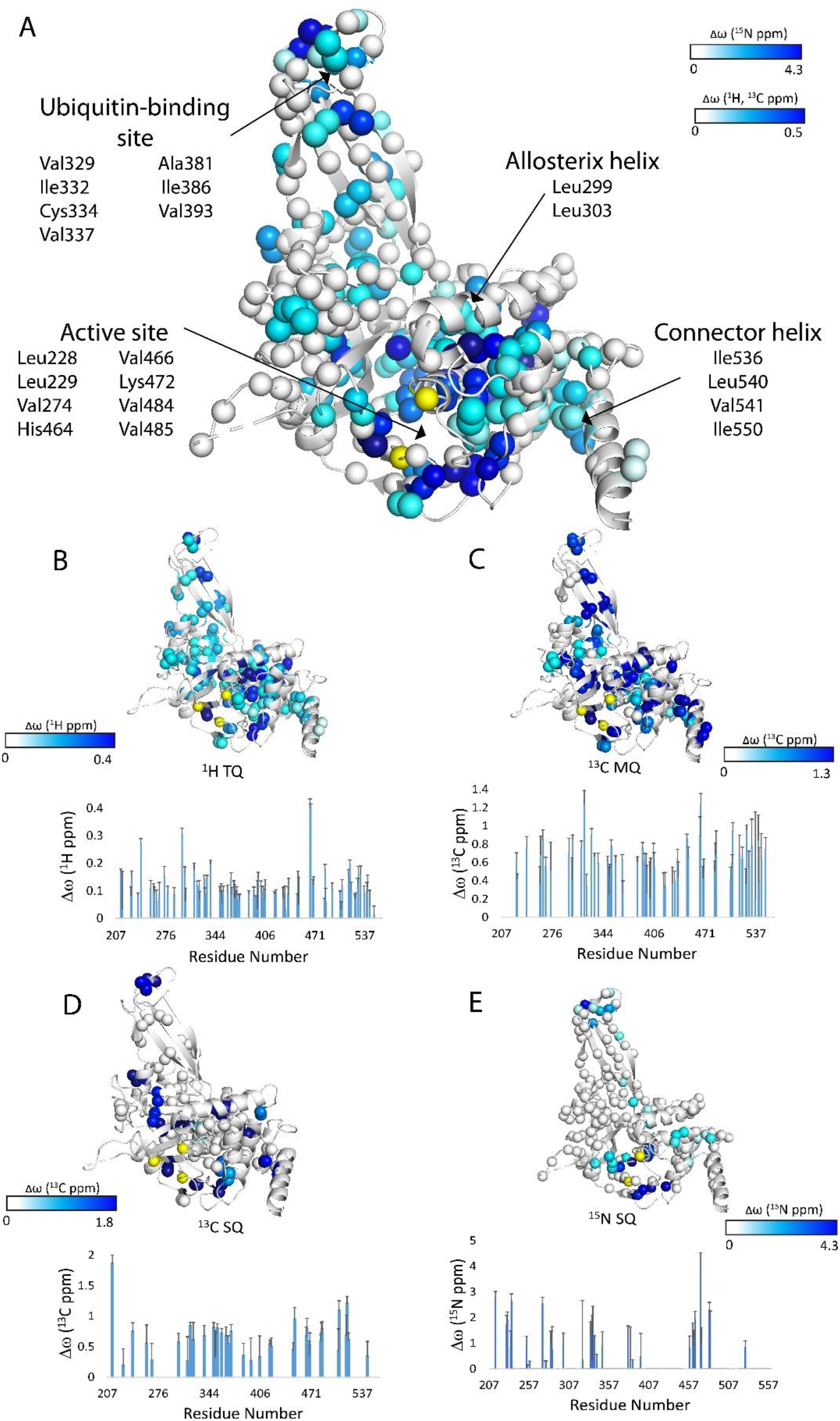
Conformationally dynamic regions of USP7. Per-residue chemical shift differences, Δω, between the exchanging major and minor conformations are plotted onto the structure of USP7-CD (PDB ID: 4m5w). Individual nitrogen atoms of the backbone amide groups and carbon atoms of the ILV methyl groups for which CPMG data were recorded are shown as spheres and colored according to their Δω values from white (smallest) to dark blue (largest). Active site triad residues (Cys223, His464 and Asp481) are depicted as yellow spheres. (**A**) Combined ^15^N backbone and ^13^CH_3_ methyl Δω. (**B**) Individual ^1^H Δω obtained from TQ CPMG. (**C**) Δω obtained from^133C-3^H MQ CPMG. (**D**) Δω obtained from ^13^C SQ CPMG. (**E**) Δω obtained from ^15^N SQ TROSY CPMG.

The ^15^N relaxation dispersions were detected for Leu229, Val274, Cys334, Val337, Ala381, His464, Val466, and Lys472 (**Figure 4E**) and clearly localized to the distinct active site and ubiquitin-binding regions of USP7. The methyl experiments also detected conformational exchange within the similar region, as well as within the allosteric helix (Leu299, Leu303) (**Figure 4B-D**), which immediately follows the SL (**Figure 1C**). Significant dispersions were detected for methyl groups of Val274, Ile332, Val337, and Val466 from the ^1^H TQ experiments (**Figure 4B**), Leu 228, Leu229, Val329, Ile332, Val337, Ile386, Val393, Val466, Val484 and Val485 from the ^13^C-^1^H MQ experiments (**Figure 4C**), and Val337, Leu386, Val466, Val485 from the ^13^C SQ experiments (**Figure 4D**). The methyl proton and carbon Δω obtained from independent SQ, MQ and TQ experiments are consistent and complementary as seen in **Figure 4A**, where these values are combined.

Despite probing different nuclei and often different residues, both, methyl and backbone CPMG consistently detected conformational exchange in the vicinity of the active site, the allosteric helix and the ubiquitin-binding fingers (**Figure 4A**), all functionally important regions that undergo rearrangement upon USP7 activation. The largest conformational changes were detected for methyl groups of Val466 (γ1 and γ2) and Leu299. Remarkably, Val466 is located only ∼4-5 Ǻ away from the His464 of the catalytic triad and expected to be most sensitive to the active site rearrangements. The Leu299 is located on the allosteric helix and has been implicated in allosteric regulation of USP7 activity [27]. Taken together, these results strongly suggest that the dynamics observed here reflect the inactive-to-active conformational transition of the USP7 catalytic domain (**Figure 2**).

### Activating mutant of USP7 slows down its conformational dynamics

After establishing that USP7-CD exists in equilibrium between its minor active and major inactive conformations we next attempted to shift this equilibrium using previously reported USP7 mutants. To shift equilibrium towards the inactive conformation, we used the C223A “de-activating” mutant, which results in a catalytically dead enzyme due to the lack of essential catalytic cysteine residue [13, 28]. To shift the equilibrium towards the active conformation, we used USP7-CD L299A “activating” mutant, which perturbs the allosteric site and was reported to increase the enzymatic activity of USP7 [27].

To characterize effect of these mutations on USP7 dynamics we have compared their^1^H TQ CMPG RDs to those of the WT USP7. Interestingly, dynamics of USP7-CD C223A remained nearly identical to the wild type, with p(B) = 5.4 ± 0.6% and k_ex_ = 8,984 ± 1,112 s^-1^ (**Table 1**). Furthermore, many of the same residues demonstrated significant Δω, including residues within the ubiquitin-binding fingers (Val337), the active site (Val466), and the allosteric helix (Leu229 and Leu303). These data demonstrate that C223A mutation prevents catalysis due to loss of the nucleophile but does not alter the dynamics of the enzyme as the mutant still transiently samples active-like conformation at nearly equal populations and rates compared to wild type.

Although L299A mutation was previously reported to cause 6-fold enhancement of the USP7 activity [27], our comparative DUB enzymatic assays showed a more modest ∼1.3-fold increase in k_cat_ (**Figure 5**). Nonetheless, L299A mutation perturbed activation dynamics of USP7 (**Table 1**). Both population of the active conformation and the exchange rate constant were affected, with p_(B)_ = 3.5 ± 0.2% and k_ex_ = 4,551 ± 773 s^-1^. While somewhat surprisingly the population of the active state has decreased, the exchange between the two states has slowed down. The resulting 1.7 times longer residence time for the active conformer in L299A compared to the WT is in agreement with 1.3-fold activity increase observed for the mutant. Similar to the WT, the regions of L299A undergoing conformational exchange included residues of the active site (Val466), the ubiquitin-binding fingers (Val337), and the allosteric helix (Leu304).

**Figure 5:**
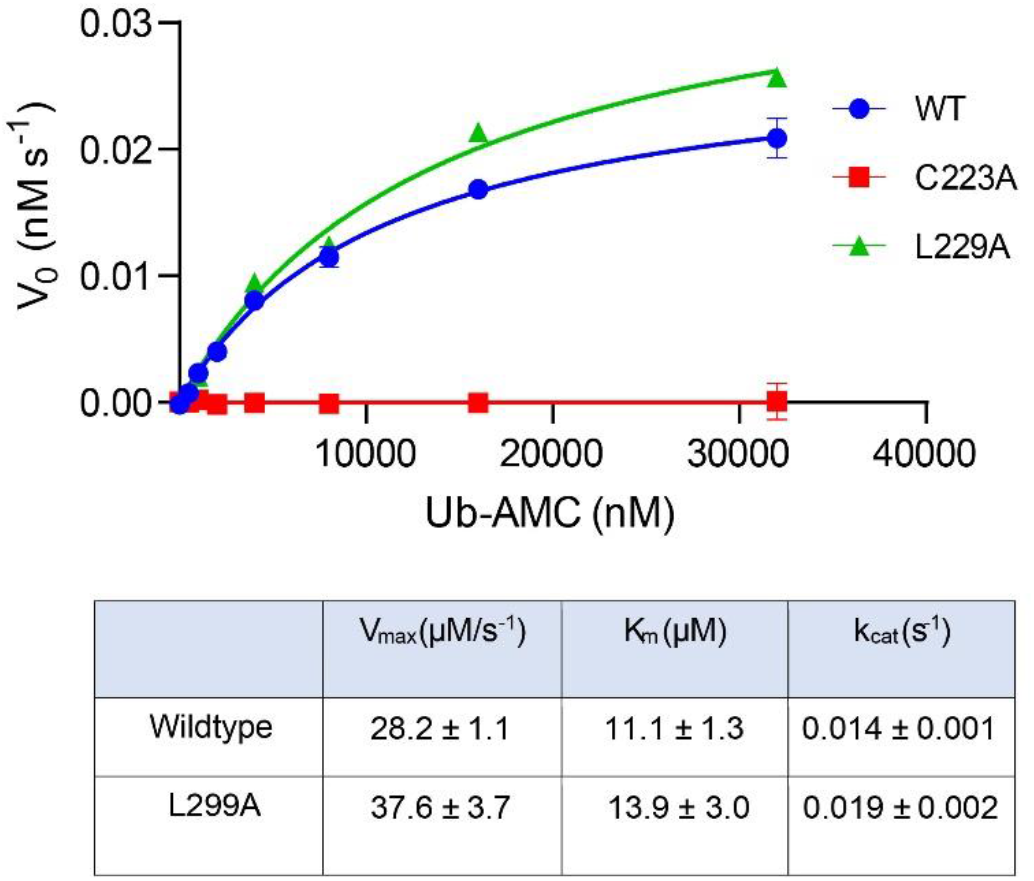
Deubiquitination enzyme kinetics assays of USP7 CD and its mutants. A plot of the initial velocities of deubiquitination reaction (V_0_) as a function of the substrate concentration (top panel), and the summary of the Michaelis-Menten analysis (lower panel).

These data suggest that inactivating C223A mutation does not alter the activation dynamics of USP7 while the activating L229A mutation slows down the exchange between the inactive and active conformations of USP7 resulting in activation of the catalytic domain.

## DISCUSSION

USP7 catalytic domain interacts with ubiquitinated substrates and undergoes active site rearrangement during the activation. Crystallography studies of the isolated domain have captured static snapshots of its apoenzyme “inactive” and substrate-bound “active” conformations (**Figure 1**), but provide no information on the dynamic conformational transitions between these distinct structural states [9, 16-19]. Using highly specialized NMR CPMG relaxation dispersion techniques to study USP7 in solution this work has revealed the intrinsic dynamic properties of the apo-enzyme on a per-residue basis.

NMR chemical shift analysis revealed that in solution the isolated USP7-CD adopts a primarily inactive conformation (**Figure 2B**). However, our NMR CPMG experiments demonstrated that USP7-CD transiently samples the active conformation in solution, even in the absence of a substrate-binding, showcasing the intrinsically dynamic behaviour of the USP7-CD.

The largest structural movements were found to localize to residues within key functional regions of USP7-CD, including the active site and ubiquitin-binding site (**Figure 4**). Val466, which is located just two residues from active site residue His464, is one example. During catalysis, His464 deprotonates the catalytic nucleophile Cys223. These residues move approximately 6 Å between USP7-CD_I_ and USP7-CD_A_ states (**Figure 1**). Due to its immediate proximity to the catalytic triad Val466 demonstrates the largest relaxation dispersion, and serves as a sensor of the rearrangement and subsequent activation of the active site. Similarly, residues of the ubiquitin-binding fingers also demonstrate dynamic movements, seemingly pre-sampling its ubiquitin-bound conformation even in the absence of a substrate (**Figure 4**).

The SL also demonstrates significant differences between the inactive and active conformations (**Figure 1C**); however, due to its high flexibility, most NMR signals in this region are broaden beyond detection. Immediately following the SL is the allosteric helix, which is the location of the L299A “activating” mutant [27] and the target of several small molecule inhibitors of USP7 [29, 30]. Interestingly, several residues within this helix demonstrate significant dynamic behaviour (**Figure 4A-B**). Due to its proximity to the SL, the dynamics seen within the allosteric helix are likely reflective of the SL rearrangement upon substrate-binding. The methyl experiments also uniquely provided insight into dynamics within the C-terminal connector helix (**Figure 4A-D**). This helix is thought to be flexible in the context of the full-length enzyme [19]. Its flexibility is essential for enzymatic activity, as it allows the regulatory C-terminal peptide tail to make important contacts with the catalytic domain to promote ubiquitin binding and enzymatic activity [9]. Several residues on the hinge and within the helix demonstrate dynamic behaviours, including Ile536, Leu540, Val541, and Ile550 (**Figure 4A**), that are likely reflective of the USP7 full-length activation mechanism.

In summary, this work has captured activation dynamics of the USP7-CD and provided insight into the transition mechanism between inactive and transiently populated active conformations of the enzyme. While it is the first such report, the phenomenon might not be unique to USP7, as structural rearrangement of the active site has also been reported for other USPs, such as USP15, for example [31]. The observed activation dynamics may be a common feature among other members of the USP family of enzymes.

## MATERIALS AND METHODS

### USP7 protein purification

Highly deuterated, amide protonated ^15^N,^13^CH_3_-ILV (^15^N/^12^C/^2^H/^13^CH_3_-Ile(δ1)/^13^CH_3_,^12^CD_3_-Leu,Val) labeled wild type (WT), C223A and L299A USP7 catalytic domain (USP7-CD) were expressed and purified in as described elsewhere [13]. Briefly, USP7-CD (residues 207-560) pET-25a-LIC plasmid was transformed into Escherichia coli BL21(DE3) cells. A 50 mL starter culture of cells in Luria Broth (LB) with 50mg/L kanamycin was grown overnight at 32°C, and was then transferred to 1L of 100% D_2_O M9 minimum media containing ^15^NH_4_Cl and ^12^C,^2^H-glucose as the sole source of nitrogen and carbon and 50mg/L kanamycin and grown until OD_600_ of 0.9-1.2 at 37°C. One hour prior to induction of protein expression, 120 mg/L of 2-Keto-3-(methyl-d3)-butyric acid-4-^13^C sodium salt (α-ketoisovalerate) (LV) and 70 mg/L 2-ketobutyric acid-4-^13^C,3,3-d2 sodium salt (α-ketobutyrate) (I) were added to the 1L culture to express ^13^CH_3_-labeled Ile-δ1 [32] or Leu-δ and Val-γ positions [33, 34]. Protein expression was induced with 1 mM 1-thio-β-ᴅ-galactopyranoside (IPTG) for 18-20h at 20 ^°^C. Cells were harvested by centrifugation, resuspended in a 20 mM phosphate, 250 mM NaCl, 10 mM imidazole, 1 mM PMSF, pH 8 buffer, lysed by sonication, and centrifuged at 15,000 rpm for 1 hour. The supernatant was filtered and loaded to a TALON HisPur cobalt resin (Thermo Scientific). Proteins were eluted in a 20 mM phosphate buffer, 250 mM NaCl and 300 mM imidazole, pH 8, and exchanged into a 20 mM Tris, 100 mM NaCl, 10 mM dithiothreitol (DTT), pH 7.4 buffer. His-tag was removed with thrombin and amide deuterons exchanged back to protons at room temperature for one week. Proteins were then subjected to size-exclusion chromatography using a HiLoad Superdex 200 column (GE Healthcare) in a 20 mM Tris, 100 mM NaCl, 10 mM DTT, pH 7.4 buffer, which is also the buffer used for NMR experiments. Final samples of WT, C223A and L299A USP7-CD contained 1.7, 1.5 and 0.9 mM protein, respectively, and 10% D_2_O.

### DUB activity assays

Deubiquitinating activity of the WT and mutants of USP7 CD was monitored using 7-amido-4-methylcoumarin (AMC) fluorescence following cleavage of the quenched minimal substrate Ubiquitin-AMC (Ub-AMC). Experiments were performed at 25°C in running buffer containing 20mM Tris PH 8.0, 0.05% CHAPS and 10 mM BME, on Corning black 384-well plates in a volume of 25 μl per well. The data was collected in time intervals of 10 minutes for a total of 150 minutes using Molecular Devices SpectraMax M3 Fluorescence Microplate Reader at excitation and emission wavelengths of 360nm and 465nm respectively. Constant USP7 enzyme concentrations of 2nM was used to hydrolyze Ub-AMC, with concentrations of the latter ranging from 0 to 32μM. The Michaelis-Menten equation was used to retrieve steady-state kinetic parameters. The experiments were performed in triplicates for all enzymes.

### NMR spectroscopy

Previously reported NMR resonance assignments of the backbone and ILV methyl groups were used [35] (BMRB ID: 51188). All Carr-Purcell-Meiboom-Gill relaxation dispersion (CPMG RD) NMR experiments were performed on an 800 MHz (^1^H) Agilent VNMRS spectrometer equipped with a room-temperature HCN Z-gradient probe at 30°C. CPMG RD measurements for WT USP7-CD included ^15^N TROSY CPMG experiment for the backbone amide groups [23] (80 scans/FID) and ^1^H triple-quantum (TQ) [24] (80 scans/FID), ^13^C single quantum (SQ) [25] (24 scans/FID) and ^13^C multiple quantum (MQ) [22] (16 scans/FID) CPMG experiments for Ile, Val, Leu ^13^CH_3_ side-chain methyl groups. In each RD experiment, a series of ^1^H-^15^N or ^1^H ^13^C 2D correlation spectra were recorded at different delays 2τ between successive 180° pulses of CPMG sequence applied during a constant relaxation period T of 30 ms (amide ^15^N), 20 ms (methyl ^13^C SQ, ^13^C MQ) or 8 ms (methyl ^1^H TQ). In addition, a reference 2D spectrum was recorded for each RD experiment with relaxation delay T set to 0. Two duplicate points for each dispersion profile were recorded for error analysis, as previously described [36]. CPMG frequency values ν_CPMG_ = 1/(4τ) ranged from 33 to 1000 Hz in ^15^N TROSY CPMG, 100 to 1000 Hz in methyl ^13^C SQ, 50 to 1000 Hz in methyl ^13^C MQ, and 250 to 3000 Hz in methyl ^1^H TQ CPMG experiments. CPMG RD measurements for USP7-CD C223A and L299A mutants included ^1^H TQ CPMG experiment [24] performed at 80 and 96 scans/FID, constant relaxation period T of 8 and 4 ms, and ν_CPMG_ of 250-3000 Hz and 250-4000 Hz, respectively.

### NMR spectra were processed with NMRPipe

[37] and analyzed in Sparky [38], as implemented in the NMRbox platform [39], using previously reported the backbone and ILV side-chain resonance assignments [35] for the WT. The C223A and L299A USP-CD resonances were assigned based on their similarity to the WT where possible. Peak intensities from pseudo 3D datasets consisting of a series of 2D spectra recorded as a function of ν_CPMG_ were converted into the effective relaxation rates (R_2,eff_) calculated as R_2,eff_(ν_CPMG_) = -1/T*ln[I(ν_CPMG_)/I_0_], where I(ν_CPMG_) is the peak intensities in 2D spectra collected with the constant relaxation delay T at different ν_CPMG_, and I_0_ is the peak intensity in a reference 2D spectrum recorded with the delay T omitted. Errors in peak intensities (ΔI) were estimated from duplicate experiments and propagated to R_2,eff_ errors as ΔR_2,eff_(ν_CPMG_) = 1/T*ΔI/I(ν_CPMG_). A minimum ΔR_2,eff_ error of 5% was assumed [36].

### CPMG RD data analysis for WT USP7-CD

The experimental CPMG RD profiles for WT USP7-CD were least square fitted to theoretical R_2,eff_ values calculated by modeling magnetization evolution during CPMG sequence by numerical solution of Bloch-McConnell equations assuming a two-site exchange between the ground state A and a low-populated state B using an in-house software cpmg_fit [40]. The data were analyzed to extract conformational exchange parameters, including the exchange rate constant (k_ex_ = k_AB_+k_BA_), population of the minor state p_B_ = 1-p_A_, and the absolute value of frequency difference between the exchanging stats (|Δω_AB_|). Initial inspection ^15^N TROSY (SQ), ^13^C SQ and ^13^C MQ CPMG profiles (showing exchange contributions to transverse relaxation R_ex_ up to 10-15 s^-1^) revealed conformational exchange in the fast on the NMR time-scale regime (|Δω| << k_ex_), in which the only exchange parameters that can be extracted from CPMG RD profiles are k_ex_ and a product p_A_p_B_Δω^2^ [41, 42]. In contrast, ^1^H TQ CPMG dispersion data report on Δω_TQ_ = 3Δω_H_, where Δω_H_ is the frequency difference between states for ^1^H SQ coherence, resulting in R_ex_ up to 9 times greater than that in a corresponding SQ CPMG data (up to ∼100 s^-1^ for USP7-CD). The significant increase in Δω_TQ_ in ^1^H TQ CPMG relative to SQ CPMG experiments effectively shifts NMR time-scale of the exchange process towards the intermediate regime (|Δω| ∼ k_ex_), in which all parameters of a 2-state exchange (p_B_, k_ex_ and |Δω|) can be obtained at the same time [41, 42]. Furthermore, relatively short ^1^H 180° pulses allow to access higher CPMG frequencies ν_CPMG_ in ^1^H TQ than in ^15^N and ^13^C CPMG experiments, providing more effective suppression of R_ex_. Therefore, despite a relatively fast relaxation of the ^1^H TQ coherence, the ^1^H TQ CPMG experiment provided in the most informative data set that allowed us to extract all 2-state exchange parameters, including the values of p_B_ and k_ex_ that were subsequently used in the analysis of ^15^N TROSY, ^13^C SQ and ^13^C MQ CPMG dispersion profiles.

^1^HTQ CPMG data for 41/84 methyl groups of WT USP7-CD (Leu228, Leu229, Leu238, Val242, Leu299, Leu260, Val316, Ile320, Val329, Ile332, Val337, Ile350, Ile360, Val370, Leu373, Leu386, Val393, Leu396, Leu402, Leu406, Ile419, Ile421, Leu437, Leu450, Val466, Leu469, Val484, Ile494, Leu505, Val507, Val517, Ile519, Leu524, Val531, Ile536) were globally fitted to a 2-state model to extract conformational exchange parameters p_B_ = 4.8±0.6% and k_ex_ = (7.8±0.5)*10^3^ s^-1^ common for all methyl groups, and ^1^H |Δω| values specific for each methyl group. To obtain ^1^H |Δω| for the remaining 43 methyl groups, ^1^H TQ CPMG profiles were then fitted using p_B_ and k_ex_ fixed to the previously determined values. In a similar manner, ^15^N TROSY, ^13^C SQ and ^13^C MQ CPMG dispersion profiles were fitted with fixed p_B_ and k_ex_ to extract frequency differences between states |Δω| for the backbone ^15^N and ILV side-chain methyl ^13^C nuclei. To obtain methyl ^13^C |Δω|, ^13^C SQ and MQ CPMG profiles for each methyl group were fitted separately, and the averaged |Δω| values obtained from the two data sets were reported.

### CPMG RD data analysis for USP7-CD mutants

To assess the effect of C223A and L299A mutations on conformational dynamics of USP7-CD, we analyzed their ^1^H TQ CPMG profiles. Noting that when approaching fast exchange regime, p_B_ and |Δω| are correlated, |Δω| values were fixed to those obtained from ^1^H TQ CPMG data fit for the WT USP7-CD. For USP7-CD C223A, the global data fit for 21/83 methyl groups (Leu229, Val242, Leu260, Leu303, Ile320, Val329, Ile332, Val337, Ile350, Ile354, Ile360, Val370, Val393, Leu402, Val466, Leu469, Leu505, Ile519, Ile550) resulted in p_B_ = 5.4±0.6% and k_ex_ = (9.0±1.1)*10^3^ s^-1^. For USP7-CD L299A, the global data fit for 19/84 methyl groups (Leu216, Leu304, Ile320, Val329, Ile332, Val337, Val370, Leu373, Val393, Leu402, Ile449, Val466, Val484, Leu505, Leu524, Ile536) resulted in p_B_ = 3.5±0.2% and k_ex_ = (4.6±0.8)*10^3^ s^-1^. This analysis made use of an assumption that C223A and L299A mutations, while shifting conformational equilibrium of USP7-CD, have only moderate effect on chemical shift differences between the exchanging states.

## AUTHOR CONTRIBUTIONS

GJV, IB and DMK conceived the research. DK performed NMR experiments. GJV processed and analyzed all data under the oversight of IB and DMK. NJ performed DUB Ubiquitin-AMC enzymatic assays. GJV, IB and DMK wrote the manuscript.

## ACKNOWLEDGEMENT

We thank Dr. Lewis E. Kay for the development of the NMR pulse sequences used in this work.

